# *Schistosoma mansoni* Granulin binds the human neutrophil receptor CD177 and modulates neutrophil activation

**DOI:** 10.64898/2026.06.27.734962

**Authors:** Martin Majer, Kelly Lee, Nicole Müller-Sienerth, Cécile Crosnier

## Abstract

To establish chronic infection in the vasculature of their infected host, schistosomes have developed multifaceted strategies of immune subversion. Extracellular parasite proteins are believed to play immunomodulatory functions, but their mode of action remains largely elusive. To investigate whether proteins secreted by the *Schistosoma mansoni* parasite have the potential to directly interact with host immune receptors, we performed a large-scale protein:protein interaction study between selected parasite proteins sharing structural similarities with known host immune effectors and a protein array of over 750 full-length human ectodomains mostly expressed by immune cells. We identified CD177 as a neutrophil receptor for *S. mansoni* Granulin (SmGrn). SmGrn exclusively bound the surface of CD177^+^ human neutrophils and led to cellular hyporesponsiveness following stimulation with LPS as evidenced by decreases in surface markers of activation, delayed reactive oxygen species production and reduced IL-8 release. In addition, human neutrophils exposed to SmGrn showed delayed apoptosis and morphological changes compatible with a more quiescent state as well as transcriptional upregulation of negative regulators of interferon signalling. These data suggest that SmGrn dampens human neutrophil response to stimulation and may lead to suboptimal function during schistosome infection.

## Introduction

Schistosomiasis is a neglected tropical disease of global health and socio-economic significance in subtropical areas, where it is endemic. In humans, the agents responsible for the disease are three main species of parasitic worms of the genus *Schistosoma* whose life cycle relies on the infection of aquatic snails and humans as intermediate and definitive hosts, respectively [1]. Human infection occurs after contact with contaminated water containing cercariae, a free-swimming infective form of the parasite. Following penetration of the skin, the parasites become schistosomula that gradually develop over the next five weeks into adult worms. To establish chronic infection, schistosomules must enter the cardiovascular system and migrate first through the lungs and then to the liver of their host where adult male and female worms pair up to begin egg-laying before reaching their final place of residence. While species responsible for the intestinal form of schistosomiasis (*S. mansoni* and *S. japonicum*) reside in blood vessels of the intestine, *S. haematobium* inhabit the venules of the bladder and cause the urogenital form of the disease. Although most of the eggs are released in the faeces (for *S. mansoni* and *S. japonicum*) or urine (for *S. haematobium*) of infected individuals, a significant proportion remains trapped in host tissues where they trigger a strong Th2 immune response that ultimately results in the formation of granulomas and the onset of the clinical symptoms of schistosomiasis [2].

One remarkable feature of parasitic helminths is their ability to establish long-lasting chronic infections by avoiding destruction by the host immune system [3]. In the absence of treatment with praziquantel, schistosomes can survive in their host’s blood vessels for several decades despite getting constantly exposed to host immune effectors [4]. Multiple mechanisms of protection have been invoked including masking of the parasite by recruitment of host molecules to its surface [5], secretion of potent parasite proteases that degrade host proteins [6-9], and the production of immunomodulatory molecules able to interfere with host cellular responses [10-14]. This downregulation of host immune function can in turn affect response to vaccination [15, 16] and co-infections [17]. Identifying new parasite antigens able to modulate immune response could not only uncover new vaccine targets (considering the paucity of suitable vaccine candidates against schistosomiasis [18]) but also reveal new molecular tools for the treatment of inflammatory conditions such as allergic and auto-immune diseases [19]. Proteomics [20-26] and transcriptomics studies [27, 28] have identified *S. mansoni* proteins expressed during the intramammalian stages, some of which are orthologous to human proteins that play important roles in immune regulation. To determine whether these parasite proteins could interfere with host immune function, we used an array of over 700 human recombinant extracellular proteins primarily expressed by immune cells and tested binding of 12 recombinant extracellular *S. mansoni* proteins [29] using SAVEXIS, a large-scale assay for the detection of direct low-affinity interactions between extracellular proteins [30]. We identified the neutrophil receptor CD177 as a binding partner for *S. mansoni* Granulin (SmGrn; Smp_170550), a secreted glycoprotein with a predicted molecular mass of 122 kDa. In the developing and adult parasite, *SmGrn* gene expression is upregulated from three hours post infection [31] and is mostly expressed by tegumental progenitors and stem cells [32]. We demonstrated that SmGrn could directly bind through its N-terminal domain to the surface of human neutrophils, and that pre-incubation of human neutrophils with SmGrn could dampen their subsequent activation by inflammatory stimuli. This interaction may have important consequences on neutrophil function and signalling.

## Material and methods

### Ethic statement

The Biology Ethics Committee of the University of York approved the use of blood collected from healthy donors (approval CC202405). All donors provided informed consent.

### Recombinant protein production, purification and quantitation

Recombinant proteins were produced by transient transfection of suspension HEK293-E or -6E cells as described previously [33]. Briefly, pTT3-based plasmids encoding the ectodomain of human secreted, GPI-anchored, or single-pass transmembrane proteins, as well as a rat Cd4 d3+4 tag, a biotinylation sequence for the biotin ligase BirA and a 6-histidine tag were transfected into HEK293 cells (at a cell density of 1-2 x 10^6^ cells/mL) as previously described [30] using a 1:3 DNA:transfection reagent mass to mass (m/m) ratio. In the case of extracellular parasite proteins, a similar approach was followed except that the endogenous signal peptide was removed and replaced by an exogenous signal peptide from the murine V_k7-33_ immunoglobulin light chain to boost recombinant protein secretion [29]. To produce enzymatically monobiotinylated proteins, each construct was co-transfected with an expression plasmid encoding the BirA biotin ligase at a 10:1 m/m ratio (e.g. 25 μg plasmid encoding the protein of interest, 2.5 μg plasmid encoding BirA and 75 μg transfection reagent for a 25 mL transfection).

Cell supernatants containing the recombinant proteins were harvested at five days post-transfection and filtered. Each supernatant was supplemented with 400 mM NaCl and 20 mM imidazole and the recombinant proteins were purified by nickel-affinity purification using a purpose-built pneumatic platform [34] and His Multitrap purification plates (Cytiva) following the manufacturer’s instructions. Following purification, the integrity and purity of recombinant proteins was assessed by SDS-PAGE and quantification of proteins was performed by Bradford assay as described previously [30]. Proteins used for cell staining and cellular assays were expressed in essentially the same way except that the expression vector for SmGrn did not contain a rat Cd4 d3+4 tag; mock transfection was performed using salmon sperm DNA and purifications were performed on HisTrap HP columns (Cytiva) using an ÄKTA Pure instrument (Cytiva).

For domain mapping experiments, deletion constructs were generated at the following residues: for CD177, Ly6 domains were expressed in pairs (CD177 d1+2; Met1-Phe208 and CD177 d3+4; Leu209-Gly408) while SmGrn was divided into 5 subdomains corresponding to domains 1+2 (SmGrn d1+2; Asn18-Val182), 3+4 (SmGrn d3+4; Ser183-Ile433), 5 (SmGrn d5; Ser434-Ile518), 6+7 (SmGrn d6+7; Ser519-Ile731) and 8 + carboxy-terminus of the protein (SmGrn d8+Cter; Ser732-Tyr920).

### SAVEXIS

Host:pathogen interactions were identified by SAVEXIS as previously described [30]. Monobiotinylated human recombinant proteins corresponding to the ectodomains of leukocyte, platelet and erythrocyte proteins were normalised to 4.5 nM in HEPES buffer saline (HBS) + 2% Bovine Serum Albumin (HBS/2% BSA) and immobilised as baits in individual wells of streptavidin-coated 384-well plates for 60 minutes before washes in HBS + 0.8 μM desthiobiotin to occupy the free streptavidin sites on the plates. Meanwhile, recombinant biotinylated proteins from the *S. mansoni* parasite were normalised to 17.5 nM in HBS/2% BSA and incubated with 10 nM horseradish peroxidase-conjugated streptavidin for 60 minutes to allow for the formation of tetramerised preys. The preys were diluted 20-fold in HBS/2% BSA before adding to all wells of the bait array. Following a 60-minute incubation, plates were washed in HBS + 0.8 μM desthiobiotin to discard unbound prey and TMB-E added to reveal horseradish peroxidase activity. Absorbance measurements at 650 nm were taken on a Tecan Spark plate reader. Positive signals were confirmed by visual inspection of the screening plates – where no signals were observed compared to the plate background (Smp_090100, Smp_210500 and Smp_132480), or parasite proteins showed broad binding to >15 human receptors (Smp_187140 and Smp_194830), no further analysis was performed.

Blocking of the SmGrn:CD177 interaction by the anti-CD177 MEM-166 antibody (Antibodies.com) was demonstrated by a competition SAVEXIS whereby the SmGrn prey was incubated with different concentrations of MEM-166 and the mixture added to the CD177 bait. The MOPC-21 isotype match (R&D Systems) was used as a negative control.

### Surface plasmon resonance (SPR)

To determine the biophysical parameters of the interaction, we used a Biacore T200 instrument (Cytiva). The ectodomain of CD177 was immobilised in the query flow cell of a streptavidin-coated SA chip (Cytiva) while the rat Cd4 d3+4 was used as the negative control in the reference flow cell. SmGrn was used as a non-biotinylated analyte. To remove protein aggregates that could affect the SPR measurements, purified SmGrn was resolved by size exclusion chromatography on a Superdex 200 Increase 10/300 column equilibrated in HBS buffer supplemented with 60 mM NaCl and 0.05% Tween 20, and the monomeric fractions used for SPR. For equilibrium binding analysis, the analyte was flowed over the surface of the chip at 37◦ C at a flow rate of 20 μL/min for 2 min until equilibrium was reached. Increasing concentrations of analyte ranging from 3.66 nM to 3.75 μM were used over successive cycles with regeneration of the chip surface with 2M NaCl at the end of each cycle. Differential binding between the experimental and reference flow cells was calculated for each concentration of analyte and an equilibrium binding constant (*K*_D_) was determined by the Biacore Analysis software. Kinetics analysis was performed in essentially the same way with a flow rate of 100 μL/min and injections of 20 seconds.

### ELISA

Monobiotinylated CD177 ectodomains (full length, d1+2 or d3+4) tagged with the rat Cd4 d3+4 tag were immobilised on a streptavidin-coated plate at a concentration of 4.5 nM in HBS/2%BSA. The proteins were incubated with 1 μg/mL of either MEM-166 (Antibodies.com) or OX68 (Novus Biologicals), two mouse monoclonal antibodies that bind human CD177 and the rat Cd4 d3+4 tag, respectively for 1hour at room temperature. After washing, the plate was incubated with a goat alkaline-phosphatase-conjugated anti-mouse immunoglobulin antibody (Sigma) at 0.2 μg/mL for one hour at room temperature. Signal was detected by colorimetric hydrolysis of a phosphatase substrate in buffer containing 10% diethanolamine and 0.5 mM MgCl_2_, pH9.2. Absorbance measurements at 405 nm were taken on a Tecan Spark plate reader.

For IL-8 measurements, undiluted supernatants from treated and stimulated neutrophils were analysed with the ELISA MAX™ Standard Set Human IL-8 kit (BioLegend), and the absorbance measured on a Tecan Spark plate reader at 450 nm.

### Neutrophil isolation and culture

Primary human neutrophils were isolated by negative selection from fresh blood of healthy donors using the EasySep™ Direct Human Neutrophil Isolation Kit (StemCell Technologies). All experiments were started within two hours of blood collection and cell isolation. The cells at a concentration of 2 x 10^6^ cells/mL were incubated in Opti-MEM I medium (Gibco) at 37°C, 5% CO_2_, treated with 100 nM untagged SmGrn or an equivalent volume of supernatant from mock-transfected cells or media alone for 15 minutes, and then activated with 1 µg/mL LPS for 15 minutes. Cells for the antibody surface staining were then collected and stained, while cells for the apoptosis assay, IL-8 ELISA and RNAseq were washed and resuspended in OPTI-MEM I containing 100 nM SmGrn or an equivalent volume of purified supernatant from mock-transfected cells or media alone, and incubated for a further four hours at 37°C, 5% CO_2_.

### Flow cytometry

Cells were incubated for 30 min at 4°C with antibodies at the following dilutions in FACS buffer (Dulbecco’s Phosphate buffered saline (DPBS) + 3% foetal bovine serum): rat BV421-conjugated anti-CD11b (clone M1/70; 1:300; BioLegend), mouse APC-conjugated anti-CD66b (clone G10F5; 1:240; BioLegend), mouse APC/Cyanine7-conjugated anti-CD177 (clone MEM-166; 1:120; BioLegend), and mouse FITC-conjugated anti-PRTN3 (clone PR3G2; 1:60, Abcam). Washed cells were resuspended in propidium iodide (PI) (1 ug/mL, BioLegend).

For detection of SmGrn binding to the cell surface, untreated and unstimulated neutrophils were incubated for 30 min at 4°C with 100 nM biotinylated SmGrn full-length, SmGrn domain 1+2, or rat Cd4 d3+4 tag as a negative control. The cells were washed and incubated for 30 min at 4°C with Streptavidin PE (1:500; BioLegend) and antibodies as described above. Washed cells were resuspended in PI as above.

For apoptosis assay, cells were stained with Annexin V FITC (1:20; BioLegend) at 4°C for 15 minutes and washed cells were resuspended in PI as above. Treatment with 2 µM staurosporine for four hours at 37°C was used to induce apoptosis and used as a positive control.

Data were acquired on a CytoFLEX LX355 (Beckman Coulter) and analysed with FlowJo (FlowJo, v. 10.0.0).

### Proteinase 3 activity assay

Freshly isolated neutrophils were seeded at 2 x 10^6^ cells/mL in DPBS into a black 96-well plate and treated with 100 nM untagged SmGrn, or an equivalent volume of purified supernatant from mock-transfected cells or media alone for 15 minutes at 37°C, 5% CO_2_, then stimulated with 1 µg/mL LPS for 15 minutes at 37°C, 5% CO_2_. The FRET substrate Abz-VAD-norV-ADRQ-EDDnp (AltaBioscience) [35] was added at a concentration of 15 µM and data acquisition started immediately on a Tecan Spark plate reader (with excitation at 320 nm and emission at 420nm) at 37°C for two hours with 45 seconds interval.

### Label-free imaging

Freshly isolated neutrophils were seeded at 2 x 10^5^ cells/mL in RPMI + 0.05% human serum albumin (HAS) in a 96-well plate with an optically clear flat bottom suitable for microscopy (Corning). The cells were treated with 100 nM untagged SmGrn, or an equivalent volume of supernatant from mock-transfected cells or media alone, and allowed to settle at 37°C, 5% CO_2_ for 30 minutes. N-Formylmethionine-leucyl-phenylalanine (fMLP; Sigma) was then added at the final concentrations 300 nM. Quantitative Phase Imaging (QPI) was performed every four minutes for three hours using a Livecyte 2 (Phase Focus) and the data analysed with the Cell Analysis Toolbox (Phase Focus, 3.12.2) to calculate optical thickness and sphericity of the cells.

### Reactive oxygen species detection

Freshly isolated neutrophils were seeded at 1 x 10^6^ cells/mL in HBSS with 10mM HEPES and 0.025% HSA into a white 96-well plate. The cells were treated with 100 nM untagged SmGrn, or an equivalent volume of supernatant from mock-transfected cells or media alone. Luminol (25 μM, Sigma) and horseradish peroxidase (1.2 U/mL, Sigma) were added simultaneously, and the mixture was incubated at 37°C, 5% CO2 for 20 minutes. Following incubation, the LPS was added at the final concentration 0.3 μg/mL and the luminescence was measured every 90 seconds for three hours using the BMG CLARIOstar plate reader at 37°C, 5% CO2. The reactive oxygen species (ROS) production was quantified as the area under the curve (AUC) and time of maximal intensity (T_max_).

### RNA sequencing

Treated neutrophils from three donors (10^5^ cells each) were washed and resuspended in Zymo DNA/RNA Shield (Zymo Research). RNAs were extracted and sequencing performed by Plasmidsaurus using Illumina Sequencing Technology and their in-house custom analysis and annotation pipeline: following fastQ generation and read-filtering, sequences were aligned to the human genome using STAR Aligner v2.7.11; quality control was performed and transcripts quantified using featureCounts. Principal component analysis was performed following raw count filtration and blind variance-stabilizing transformation (VST) using DESeq2; differential expression and functional enrichment pipelines were performed by Plasmidsaurus using edgeR v4.0.16 and GSEApy v0.12 and the MSigDB Hallmark geneset, respectively. Gene ontology and biological pathway overrepresentation test was performed by PANTHER on the 100 most differentially expressed coding transcripts using PAN-GO human functionome v2.0 [36] and the Fisher’s exact test with false discovery rate correction.

### Statistical analysis

The statistical analysis and graphs visualization were performed in GraphPad Prism (v. 10.1.2). Significance was set at P value <0.05, with the following levels: *P<0.05, **P<0.01, ***P<0.001. The statistical tests used are indicated in the figure legends.

## Results

### Schistosoma mansoni Granulin interacts with the human surface receptor CD177

We have recently used published proteomics and transcriptomics data to assemble a library of recombinant extracellular *S. mansoni* proteins expressed during the intramammalian stages of the parasite lifecycle [29, 37]. Within this library, we sought to identify secreted molecules with possible immune regulatory function by looking for orthology with known human immune proteins; we selected 12 candidates that shared over 25% amino-acid sequence similarity with human proteins involved in immune function (Table 1). To determine whether these parasite candidates could interact with human immune effectors, we used the SAVEXIS assay to interrogate an array of 753 recombinant human extracellular domains, the majority of which are expressed by leukocytes, platelets and megakaryocytes (Table S1). Three parasite proteins (Smp_090100, Smp_210500 and Smp_132480) did not show any significant binding, while another two showed broad binding to >15 human receptors (Smp_187140 and Smp_194830) and were not analysed further. For the remaining seven parasite proteins, z-scores were calculated across the whole human library and interactions with a z-score>5 were studied (Fig. 1A). Interactions were identified between *S. mansoni* fasciclin-domain containing protein and the human protein EPHB1 (Fig. 1B), *S. mansoni* Granulin (SmGrn) and the human proteins CD177 and IGFBP1 (Fig. 1C), and the *S. mansoni* hypothetical protein UPF0506 with the human protein IGFBP1 (Fig. 1D). In addition, most parasite proteins interacted with either CD209 (also known as DC-SIGN or CLEC4L) and CLEC4M (also known as L-SIGN or CD299), two C-type lectin domain-containing proteins that have shown promiscuous binding behaviour in previous protein interaction studies [30], most likely due to their binding to mannose-containing glycans present on the parasite preys (Fig. 1B-E). To validate our initial observations, we produced new protein preparations for EPHB1, CD177 and IGFBP1 as well as the twelve *S. mansoni* recombinant proteins and tested whether the interactions identified in the primary assay could be replicated and reciprocated (by using the parasite proteins as baits and the human ectodomains as preys). While all interactions could be replicated in the same orientation as the initial assay, the interaction between CD177 and SmGrn was the only one that could also be reciprocated (Fig. 1F) and will be the focus of our study. Furthermore, to determine whether the functional outcome of SmGrn interaction with neutrophils could be investigated in the mouse model, we tested whether the parasite protein could interact with mouse Cd177 (MmCd177). However, no interaction was observed with the murine receptor using SAVEXIS (Sup. Fig. 1).

**Table 1.**
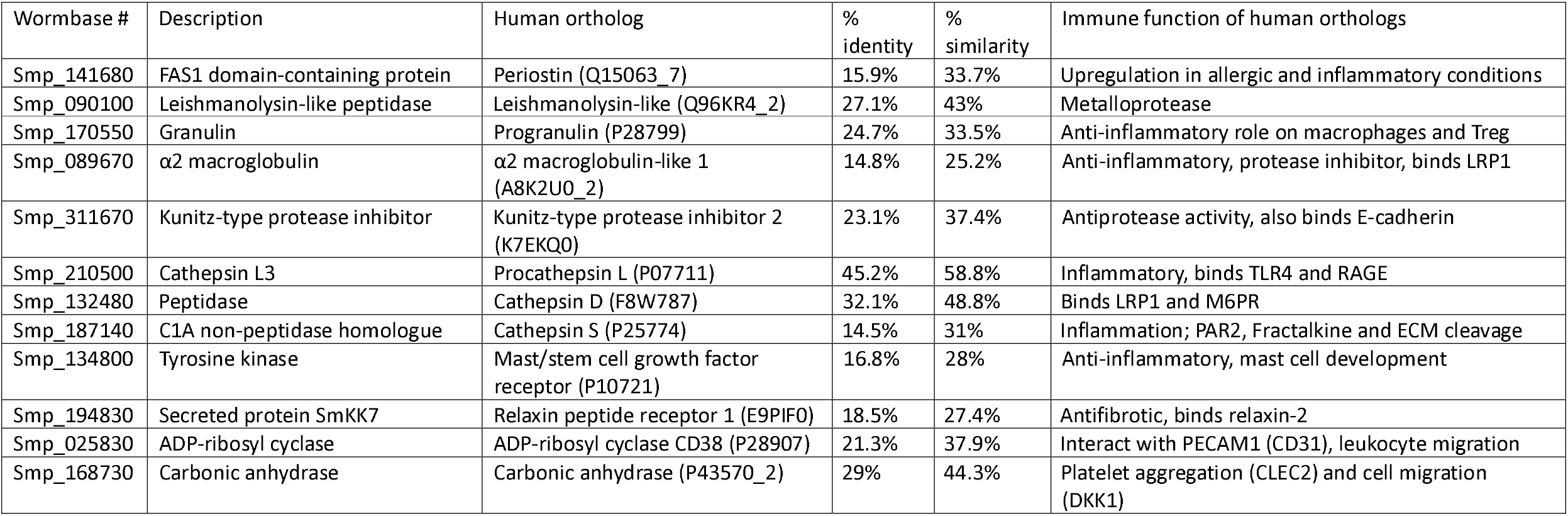
List of selected recombinant *Schistosoma mansoni* proteins whose human orthologs have immune function.

**Figure 1.**
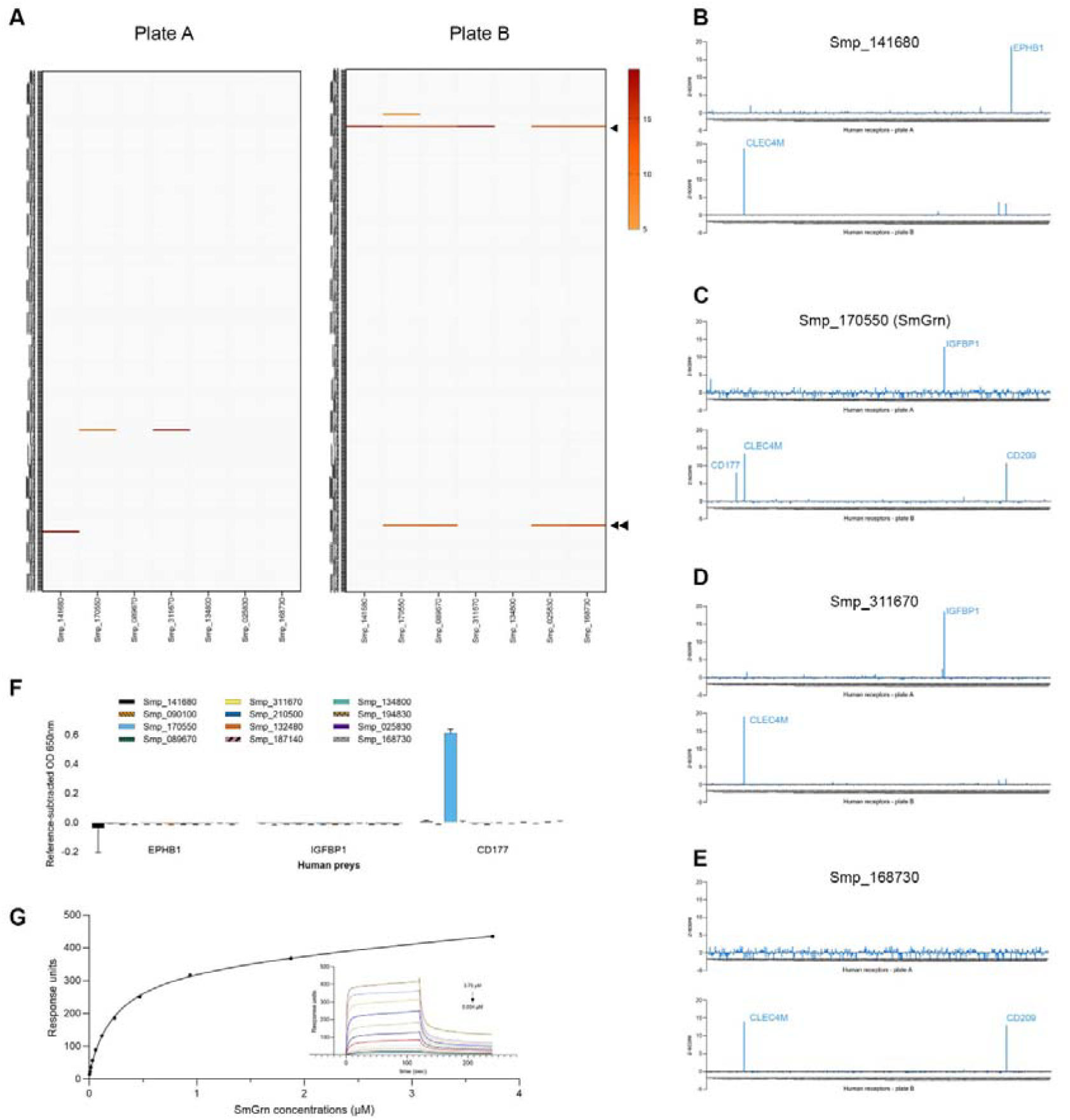
*S. mansoni* Granulin interacts with the human receptor CD177. **A**. Heatmap showing all the interactions with a z-score greater than 5 from the primary SAVEXIS assay. Five parasite proteins that showed no signal or interacted with more than 15 human receptors were excluded from the heatmap. Human proteins were arrayed as baits over two 384-well plates: plate A (left) and B (right). Each human protein is presented in a row while parasite preys are arranged in columns. The promiscuous binders CLEC4M and CD209 are indicated on plate B as single and double arrowheads, respectively. **B-E**. Individual binding profiles over plate A (top) and plate B (bottom) for Smp_141680 (B), Smp_170550 (C), Smp_311670 (D) and Smp_168730 (E). Note that Smp_089670, Smp_134800 and Smp_025830 showed similar binding profile as the one shown in E. **F**. SAVEXIS reciprocation: the twelve selected parasite proteins were arrayed as baits and tested for interaction with EPHB1, CD177 and IGFBP1. Only the interaction between CD177 and Smp_170550 (SmGrn) could be reciprocated. Reference subtracted data; error bar = SD from n=3 technical replicates G. Equilibrium binding analysis of SmGrm used as an analyte over tagged human CD177 immobilised as a ligand on a streptavidin-coated chip. The tag only control was used in the reference flow cell.

To characterise the biophysical parameters of the interaction, we used surface plasmon resonance. Increasing concentrations of SmGrn analyte were flowed over successive cycles on the chip surface containing CD177 until binding equilibrium was reached. Equilibrium binding analysis demonstrated direct, saturable binding of SmGrn to CD177 with K_D_ = 0.37 ± 0.03 μM (Fig. 1G). In parallel, kinetics analysis assuming a 1:1 binding model produced an association rate constant k_a_ = (1.76 ± 0.08) x 10 M^-1^s^-1^ and a dissociation rate constant k_d_ = 0.22 ± 0.01 s^-1^ resulting in a K_D calc_ = 1.2 μM, in good agreement with the equilibrium binding analysis (Sup. Fig. 2).

### SmGrn and CD177 interact through their amino-terminal domains

The extracellular domain of human CD177 is composed of four Ly6 domains organised in pairs that form two tight globular subdomains [38], while SmGrn is a large secreted protein composed of 8 full granulin domains, a half-domain located between domains 2 and 3 and an incomplete C-terminal domain missing two of the canonical cysteine residues. To identify which regions of the proteins were involved in this interaction, we generated deletion constructs for both CD177 and SmGrn: for CD177, each pair of Ly6 domains was expressed individually while SmGrn was divided into 5 subdomains (Fig. 2A). Using the SAVEXIS assay, we showed that the N-terminal subdomain of CD177 (CD177 d1+2) interacted with the amino-terminal end of SmGrn (SmGrn d1+2) (Fig. 2B). This interaction produced a stronger binding signal than full-length SmGrn, possibly due to better accessibility of SmGrn d1+2 to the human receptor. To further demonstrate specificity of the binding, we used MEM-166, a monoclonal antibody directed against the amino-terminal subdomain of CD177 (Sup. Fig. 3). When co-incubated with a SmGrn prey, MEM-166 could outcompete the prey for binding to CD177 in a dose-dependent and saturable manner (Fig. 2C) at concentrations as low as 1.25 μg/mL (~8 nM).

**Figure 2.**
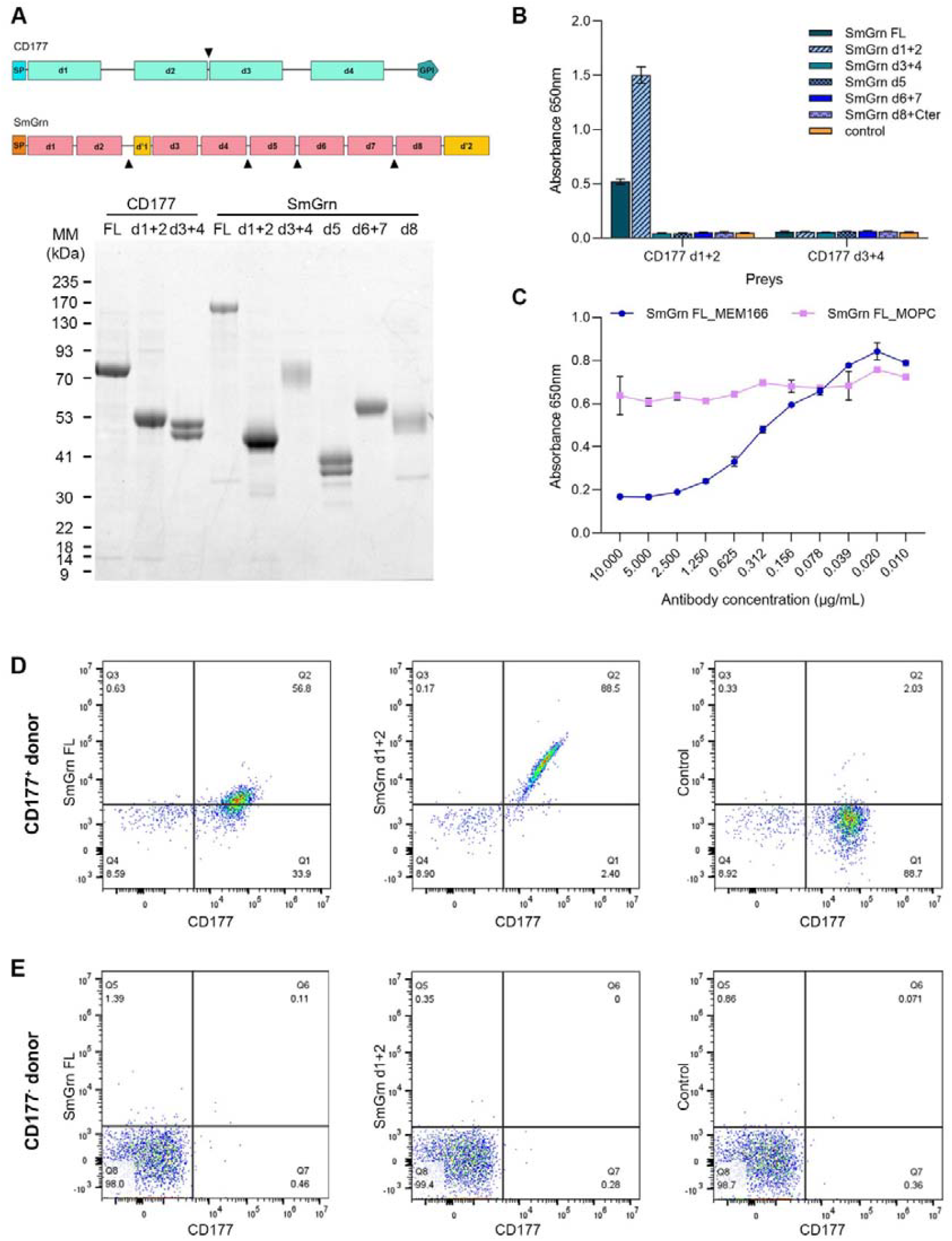
*S. mansoni* Granulin and human CD177 bind each other through their N-terminal domain. **A**. Schematic representation of CD177 and SmGrn (top) and protein subdomains used for the mapping of the interaction (bottom). Both proteins contain a signal peptide for secretion (SP). CD177 is composed of four Ly6 domains (d1-d4) and is anchored to the neutrophil surface through a glycosylphosphatidylinositol (GPI) structure, while SmGrn is composed of 8 granulin domains (d1-d8) and two incomplete domains (d’1 and d’2). Black arrowheads indicate the position of the truncation sites to produce the different subdomains. The purified subdomains were resolved on a 4-12% polyacrylamide gel and compared to the full-length (FL) ectodomains of CD177 and SmGrn. **B**. SAVEXIS assay demonstrating interaction between the N-terminal domains of CD177 (CD177 d1+2) and SmGrn (SmGrn d1+2). The different subdomains of SmGrn or the Cd4 tag alone (control) were arrayed as baits on a microtitre plate and tested for binding to the N-terminal and C-terminal subdomains of CD177 (CD177 d1+2 and CD177 d3+4, respectively) used as preys. Error bar = SD from n=3 technical replicates. One representative example of 3 biological replicates. **C**. The anti-CD177^+^ antibody MEM-166 can block the interaction between CD177 and SmGrn. SmGrn was used as an HRP-tagged prey and mixed to varying concentration of either MEM-166 anti-CD177 antibody or MOPC-21 isotype control; the mixtures were then transferred to a microtitre plate containing the CD177 bait, and interactions detected by colorimetric turnover of an HRP substrate. Error bar = SD from n=3 technical replicates. One representative example of 2 biological replicates. **D, E**. SmGrn binds specifically to CD177^+^ neutrophils. Representative flow cytometry dot plots of blood-isolated neutrophils from a CD177^+^ donor (with 91% CD177 neutrophils) (D), and a CD177^+^ donor (E). Cells were stained with an anti-CD177 antibody and biotinylated proteins: SmGrn (left), SmGrn d1+2 (centre), or a Cd4 tag only as a control (right), and detected via streptavidin-PE.

### SmGrn binds the surface of primary human neutrophils in a CD177-dependent manner

CD177 is primarily expressed on the surface of neutrophils and exhibits a peculiar bimodal distribution whereby only a fixed fraction of neutrophils is CD177-positive (CD177^+^) in any given individual throughout their life [39, 40]. Although expression level of CD177 on the surface of activated CD177 neutrophils can increase, the relative proportion of positive neutrophils in an individual remains unchanged and ranges from 0 to 100%, with an estimated 3-5% of the general population showing no CD177^+^ expression on any of their neutrophils [41, 42].

To further characterise the interaction, we isolated human primary neutrophils from blood donors with various degrees of CD177 positivity, ranging from 0 to 91%. Following incubation with PE-conjugated SmGrn tetramers (Sup. Fig. 4), surface staining was observed with SmGrn and SmGrn d1+2 on neutrophils from CD177^+^ individuals (Fig. 2D) but not on the neutrophils of a CD177^-^ individual (Fig. 2E). In particular, SmGrn d1+2 showed a bimodal distribution that remarkably paralleled the one observed for CD177 with the proportion of SmGrn-stained cells mirroring that of CD177^+^ neutrophils (Sup. Fig. 5).

### SmGrn does not trigger neutrophil activation but modulates response to LPS stimulation

To study the effect of SmGrn on neutrophil activation, we stimulated neutrophils with lipopolysaccharides (LPS) and looked at two surface markers: the integrin ITGAM (CD11b or Mac-1, a marker of tertiary granules) and the cell adhesion molecule CEACAM8 (CD66b, a marker of secondary granules) whose surface expression increases in activated neutrophils (Fig. 3A and Sup. Fig. 6) [43]. Incubation of human CD177^+^ neutrophils with SmGrn did not affect surface expression of CD11b or CD66b in resting neutrophils (Fig. 3B and 3D, left panels). Stimulation with LPS led to an increased surface staining of both markers; however, the increase in CD66b level was dampened in neutrophils that had been pre-incubated with SmGrn, suggesting that the parasite protein may interfere with the neutrophil response to stimulation (Fig. 3C and 3D and Sup. Fig. 6).

**Figure 3.**
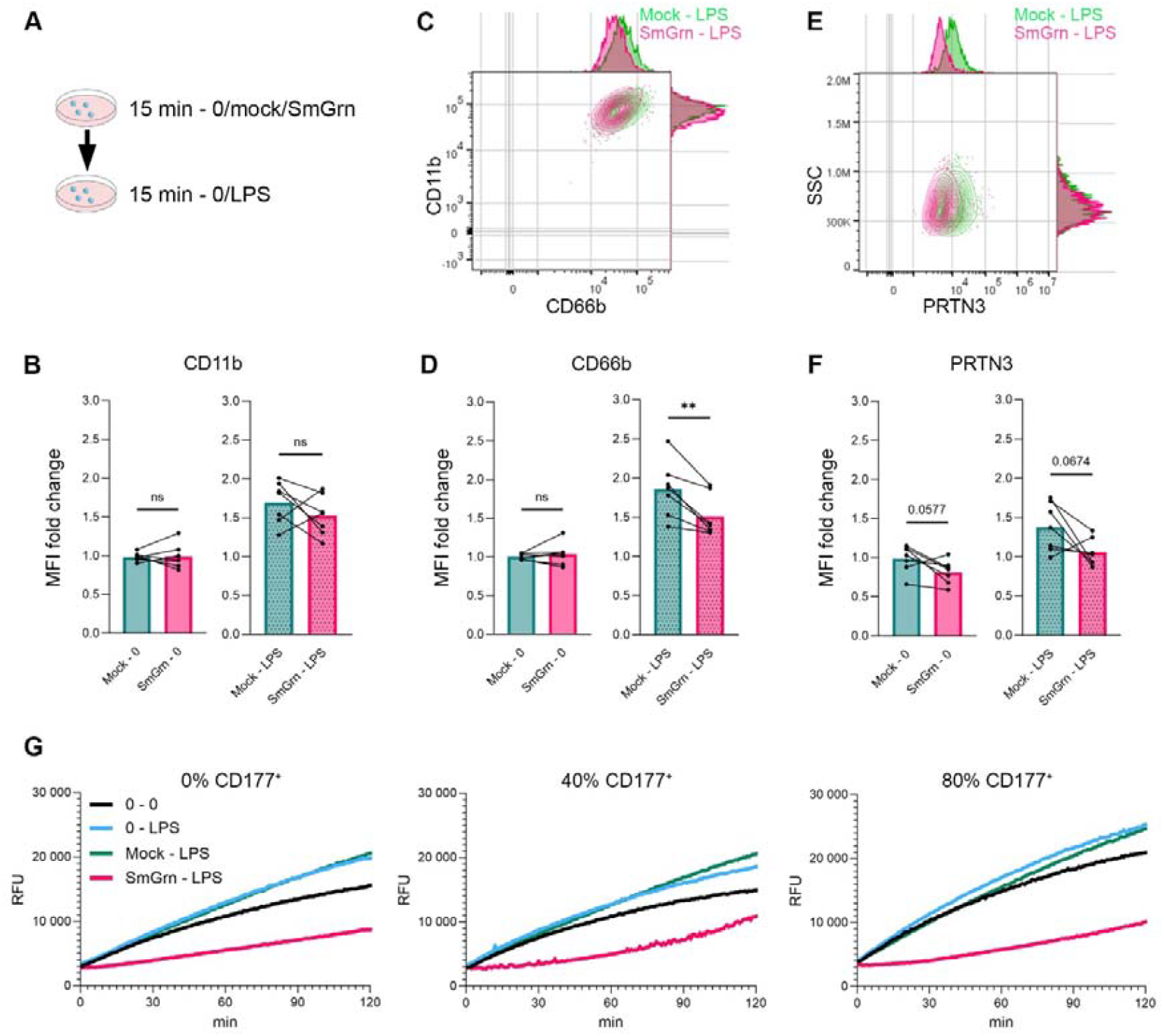
SmGrn reduces the neutrophil response to LPS stimulation. **A**. Experimental design: neutrophils were incubated for 15 minutes in the presence of either buffer (0), a mock transfectant (mock), or *S. mansoni* Granulin (SmGrn) before incubation for 15 minutes with either buffer (0) or lipopolysaccharides (LPS). **B**. Median fluorescence intensity (MFI) fold change of CD11b on neutrophils compared to untreated-unstimulated samples. **C**. Representative plot of CD11b and CD66b surface expression on neutrophils with or without pre-incubation with SmGrn. **D**. Median fluorescence intensity (MFI) fold change CD66b on neutrophils compared to untreated-unstimulated samples. **E**. Representative plot of PRTN3 surface expression on neutrophils. **F**. MFI fold change of PRTN3 on neutrophils against untreated-unstimulated samples. **G**. Enzymatic activity of PRTN3 released from neutrophils isolated from 3 donors with fractions of CD177^+^ neutrophils ranging from 0 to 80%. (0-0) correspond to untreated and unstimulated neutrophils. C, D, F. Bars represent the means, and connected dots represent individuals (n = 7 different donors from 3 independent experiments with fractions of CD177^+^ neutrophils ranging from 40% to 93%). The statistical significance was assessed by the paired t-test and indicated by ** P<0.01.

### SmGrn affects surface expression and activity of Proteinase 3

CD177 is a known receptor for Proteinase 3 (PRTN3), a serine protease mostly stored in the primary azurophilic granules of resting neutrophils [44]. Upon cell activation, PRTN3 is released from the granules and while part of the proteinase pool is discharged in the extracellular space, a fraction associates with CD177 to be displayed on the neutrophil surface [45]. We therefore investigated whether SmGrn could interfere with surface expression of PRTN3 (Fig. 3E, F). CD177^+^ resting neutrophils pre-incubated with SmGrn showed a small although not statistically significant decrease in surface PRTN3 staining (p=0.0577, Fig. 3F). Stimulation with LPS led to increased PRTN3 surface expression in CD177^+^ neutrophils preincubated with buffer or mock controls. However, when preincubated with SmGrn, PRTN3 expression remained at levels similar to that of resting neutrophils (Fig. 3F and Sup. Fig. 6).

To validate these observations, we incubated cells from three donors with no, 40% or 80% CD177^+^ neutrophils in the presence of a substrate that emits fluorescence upon cleavage of a quencher by PRTN3 [35]. Stimulation with LPS of neutrophils pre-incubated with the mock or buffer controls showed increased catalytic activity, as expected from the increased presence of PRTN3 at the cell surface. Strikingly, processing of the substrate was strongly inhibited in SmGrn-treated neutrophils (Fig. 3G). This inhibition was also observed with neutrophils from a CD177^-^ individual, suggesting that decrease in catalytic activity happened irrespective of PRTN3 association with CD177.

### SmGrn dampens the inflammatory response of neutrophils

PRTN3 present on the surface of neutrophils has recently been shown to bind the adhesion G-protein coupled receptor ADGRG3 to activate neutrophils through cleavage of the Protease-activated receptor 2 (PAR2), which in turn leads to IL-8 release from neutrophils [46]. To determine whether the decrease in PRTN3 activity observed in SmGrn-treated cells was associated with a downregulation of neutrophil activation, we measured IL-8 production in SmGrn-treated and control CD177^+^ neutrophils stimulated with LPS over a period of 4 hours (Fig. 4A). A significant reduction in IL-8 secretion was observed in neutrophils that had been pre-incubated with the parasite protein compared to controls (Fig. 4B). In addition, we measured the release of reactive oxygen species (ROS) over time following LPS stimulation. Total amount of ROS was measured by calculating the area under the curve in neutrophils pre-incubated with control or SmGrn: no significant difference was observed between the treatment conditions (Fig. 4C). However, we observed that the time of maximal ROS release (T_max_) was delayed by 15 to 20 minutes in neutrophils exposed to SmGrn compared to control (Fig. 4 D, E and Sup. Fig. 7). These combined observations could be hallmarks of more quiescent neutrophils, which tend to have a rounder shape compared to flatter, more adherent activated neutrophils [47]. To investigate whether SmGrn could influence these morphological changes, we performed label-free microscopy experiments to evaluate the optical thickness and sphericity of neutrophils. Following incubation with SmGrn or control and activation with fMLP, cells were monitored over a period of three hours. When compared to control, neutrophils incubated with SmGrn showed an increased sphericity (Fig. 4F) and optical thickness (Fig. 4G) consistent with a less activated state. Finally, we investigated whether SmGrn could affect neutrophil viability. Early and late apoptosis was measured by Annexin V and propidium iodide staining, respectively [48] on either resting or LPS-stimulated CD177^+^ neutrophils (Sup. Fig. 8). In resting neutrophils, incubation with SmGrn or control had the same effect on the relative percentage of viable and apoptotic cells compared to untreated conditions (~75% viable cells). Stimulation with LPS led to a marked increase in apoptosis. However, this effect was again dampened when neutrophils had been pre-incubated with SmGrn and led to a small increase in the fraction of viable cells (Fig. 4 H, I and Sup. Fig. 9). Together these data suggest that SmGrn help maintain human neutrophils in a hyporesponsive state after stimulation.

**Figure 4.**
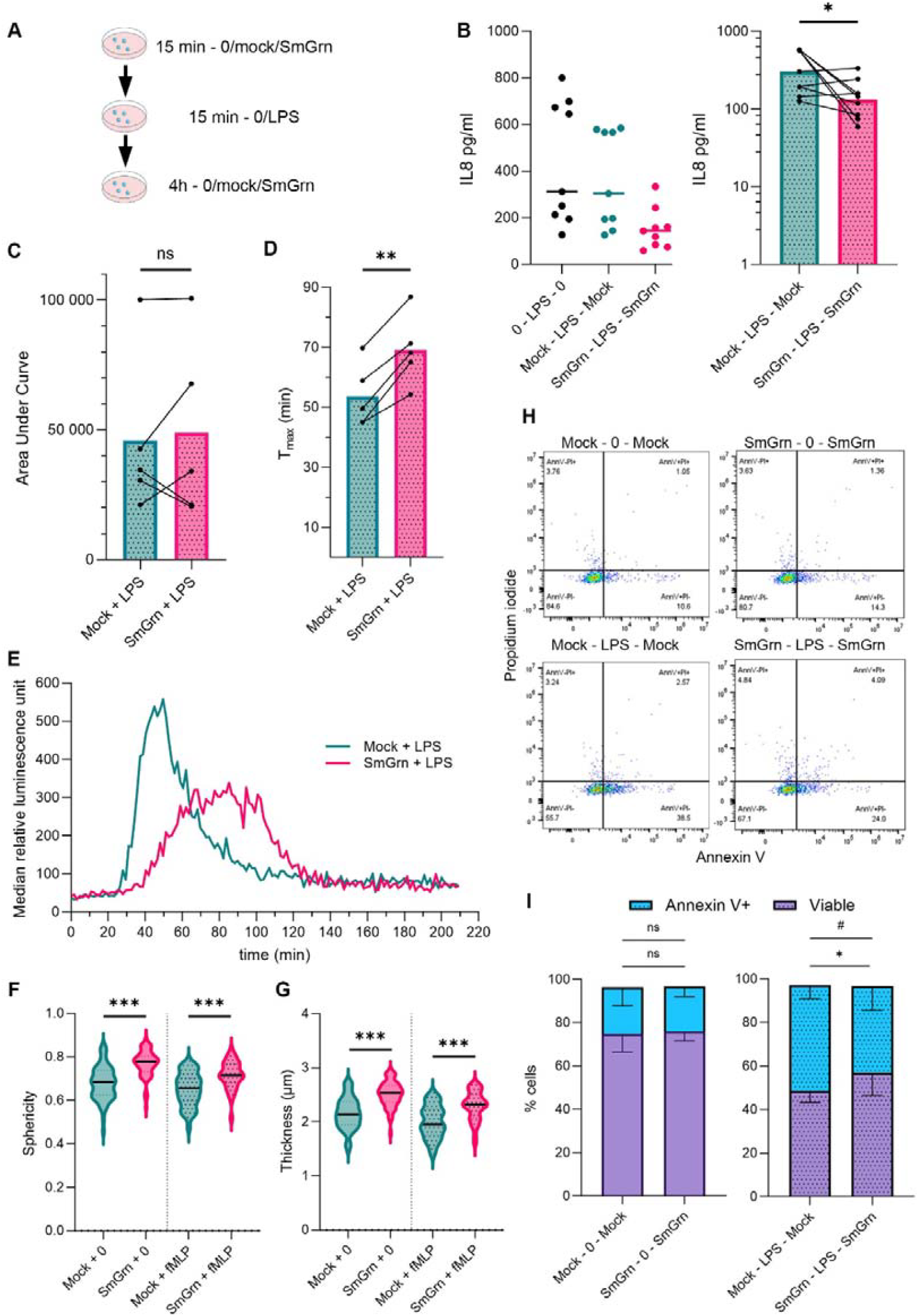
SmGrn reduces neutrophil IL-8 release and promotes neutrophil quiescence. **A**. Experimental design: neutrophils were incubated for 15 minutes in the presence of either buffer (0), a mock transfectant (mock) or *S. mansoni* Granulin (SmGrn) before incubation for 15 minutes with either buffer (0) or lipopolysaccharides (LPS) and incubation for a further four hours. Supernatants were then collected for IL-8 measurements. **B**. Raw measurements (left) and log-normalised data (right) of IL-8 released by neutrophils (n=9 individuals from 4 independent experiments with fractions of CD177^+^ neutrophils ranging from 54 % to 94%). Statistical significance was assessed by paired t-test and indicated by * P<0.05. **C**. Reactive oxygen species (ROS) production measured by area under the curve in neutrophils from five donors incubated with either mock transfectant (mock) or *S. mansoni* Granulin (SmGrn). Bars represent the mean and connected dots represent individuals’ means of technical triplicates from two independent experiments (n=5) with fractions of CD177^+^ neutrophils ranging from 45% to 93%. **D, E**. Time of maximal ROS release (T_max_) from the same donors as in C: bars (D) represent the mean and connected dots represent individuals; median luminescence over time is shown in E. **F, G**. The morphology of neutrophils was assessed by sphericity (F) and optical thickness (G). Medians from three donors with CD177^+^ neutrophil fractions ranging from 45% to 83% are shown after 60 minutes. One representative example of two biological replicates. The statistical significance was assessed by unpaired t-test and indicated by *** P<0.001. **H, I**. Apoptosis assay: representative dot plots for the different treatment conditions are shown in H. The proportions of viable and Annexin V^+^ neutrophils (I) were represented as the mean with SD (n = 5 donors from 3 independent experiments with fractions of CD177^+^ neutrophils ranging from 43% to 93%). The statistical significance was assessed by paired t-test and indicated by * P<0.05 for the viable group and # P<0.05 for the Annexin V^+^ group.

### RNA sequencing shows an upregulation of interferon-stimulated genes in neutrophils exposed to SmGrn

To further investigate the effect of SmGrn on human neutrophils, we performed RNAseq analysis on cells isolated from three different donors. Principal component analysis showed a marked effect of SmGrn as opposed to controls or untreated neutrophils (Fig. 5A). Incubation with the parasite protein led to the increased transcription of 215 genes (fold change >5; adjusted p value = 0.01; Table S2) with only two downregulated genes (*NECTIN1* and *NACC2*). Interferon-stimulated genes (ISGs) were significantly upregulated, in particular those associated with negative regulation of interferon signalling such as *IDO1* [49], *USP18* [50], *SOCS1* [51], *MT2A* [52] and *NT5C3A* [53] (Fig. 5B, C). GO term gene enrichment analysis on the 100 most upregulated genes in SmGrn-treated neutrophils compared to control (Fig. 5D) showed that biological processes such as IL27-mediated signalling pathway and negative regulation on type II interferon production were overrepresented. These observations suggest that SmGrn drives the neutrophil transcriptional profile towards an anti-inflammatory phenotype.

**Figure 5.**
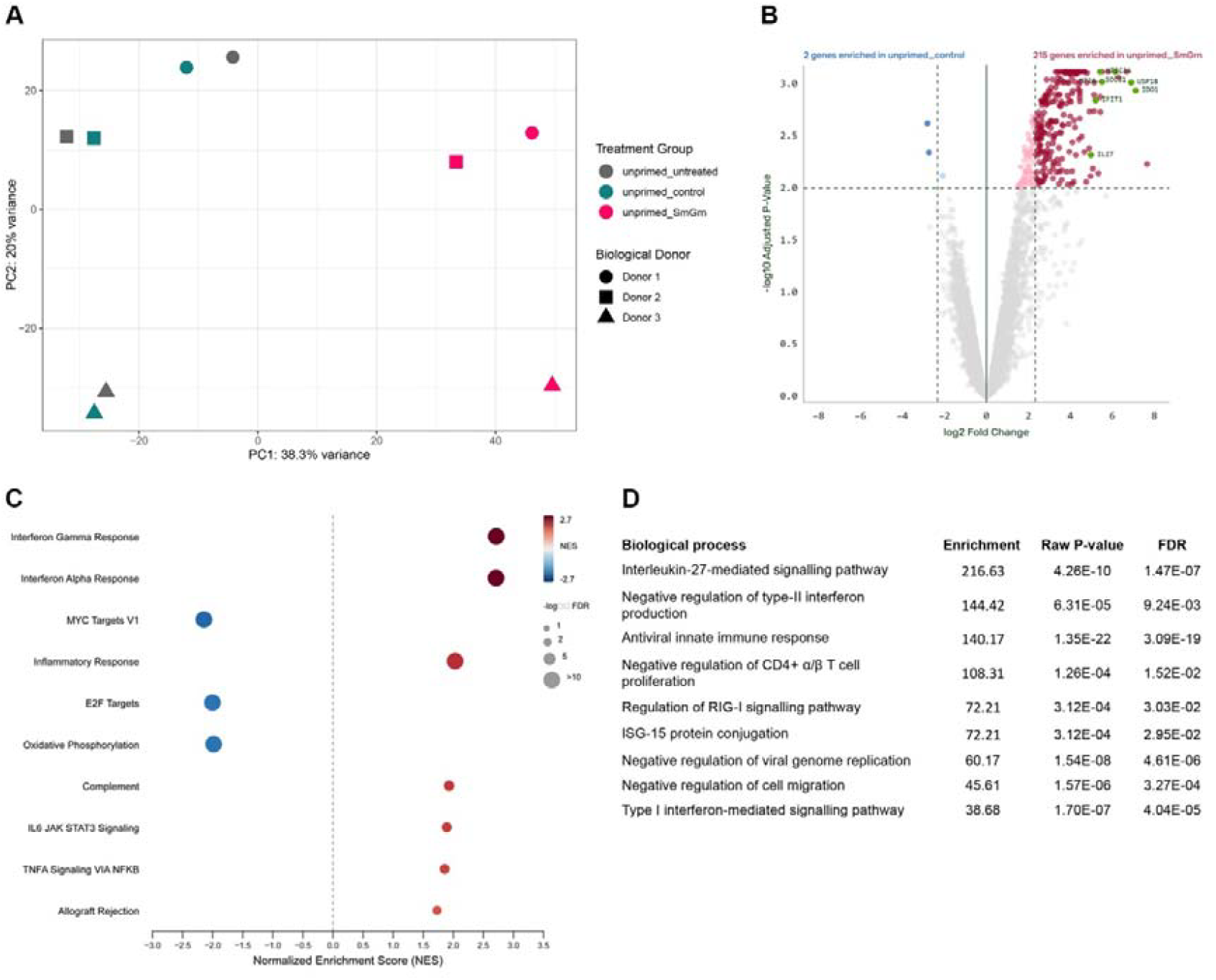
Incubation of human neutrophils with SmGrn leads to an upregulation of genes associated with negative control of interferon signalling. **A**. Principal component analysis for the three different treatment conditions (untreated, incubated with either SmGrn or mock transfectant) on human neutrophils from three donors where each dot represents a donor and each colour a different treatment: incubation with SmGrn had a substantial effect on neutrophil transcriptome. **B**. Volcano plot of genes significantly upregulated (fold change >5; adjusted p value = 0.01) in SmGrn-treated neutrophils compared to mock controls: 215 transcripts were significantly upregulated after SmGrn incubation while only two transcripts were significantly downregulated. **C**. Pathway enrichment analysis showing significant upregulation of genes associated with interferon signalling in SmGrn-treated neutrophils. **D**. Most significant biological processes identified following GO term enrichment analysis on the 100 most upregulated coding transcripts in SmGrn-treated neutrophils.

## Discussion

Despite neutrophils being considered the first line of defence against pathogens, surprisingly little is known about their role in humans during the early stages of *S. mansoni* infection as most of the functional data have been obtained from the murine model of infection [54-56]. Although killing of *S. mansoni* somules by human granulocytes has been observed in vitro, it appears to be more potently mediated by eosinophils rather than neutrophils in the presence of complement or antibodies+complement [57-59]. Co-incubation of *S. mansoni* somules with human granulocytes in the presence of either lectins, anti-somule antibodies or active complement on the parasite surface has shown endocytosis of patches of parasitic material by neutrophils without disruption of the tegumental membrane and in some instances, fusion of neutrophil and upper tegumental membranes where endocytosis has occurred [60, 61]. Interestingly, less than 5% of neutrophils showed signs of degranulation in these studies [61]. There is now growing evidence that proteins produced by the parasite can affect neutrophil function: the serine protease inhibitors SmKI-1 and SmSerpin-p46 can have potent inhibitory effect on neutrophil elastase activity [62, 63]. In addition, GPI-anchored *S. mansoni* proteins released from the parasite surface can form complexes with host lipoproteins, which negatively affect cell viability when endocytosed by neutrophils [64].

In this study, we identified CD177 as a receptor for SmGrn. CD177 is a GPI-anchored protein primarily expressed on neutrophils and to a lesser extent on breast epithelial cells [65]. Its upregulation has also been described in tumour-infiltrating regulatory T cells [66]. In human neutrophils, CD177 is stored in secondary and tertiary granules from which it can be rapidly released to the cell surface upon neutrophil activation, and its expression is characterised by a bimodal distribution whereby only a fraction of cells is CD177^+^. Being devoid of intracellular domain, CD177 triggers downstream signalling through its cis-interaction with α_M_β_2_ integrin (Mac-1, CD11b/CD18) to promote neutrophil adhesion [67]. In addition, CD177 acts as the main binding partner for PRTN3, a soluble serine protease released from primary granules. The presence of the CD177:PRTN3 complex on the surface is believed to be transient: both proteins are rapidly internalised and recycled in a process that lasts no more than 30 minutes [68]. Interaction of the CD177-tethered PRTN3 with the adhesion G-protein coupled receptor G3 (ADGRG3, GPR97) induces conformational change of PRTN3 resulting in cleavage of PAR2 and inflammatory activation of neutrophils [46].

We demonstrated that SmGrn binding to the neutrophil surface was mediated by its interaction with CD177 as evidenced by the lack of binding in CD177-negative individuals, and that this interaction was mediated by its amino-terminal domain. Expression of *SmGrn* is upregulated as early as 48 hours post-infection of the mammalian host [27] and increases further in the juvenile male and female worms [31]. Single-cell sequencing of two-day old somules has identified *SmGrn* transcripts in the precursors of the tegument, which lines the outer surface of the parasite [69], while expression in adult worms is more broadly distributed across tissues [32]. Antibody reactivity to SmGrn peptides has been identified in the sera of self-curing rhesus macaques, starting at 10 weeks post-infection and maintained thereafter [70].

Mammalian granulins exert pleiotropic functions pertaining to inflammation, cell proliferation, wound healing, and neurodegeneration. While the human precursor Progranulin has been shown to have anti-inflammatory function through its interactions with Tumour Necrosis Factor Receptor 1 and 2 (TNR1A and TNR1B) [71] and its extracellular level is regulated by Sortilin (SORT1) [72], individual granulin domains released following Progranulin proteolysis can have activating function on neutrophils [73]. Although both TNF receptors and SORT1 were amongst the human receptors represented on the array used in our study, we did not detect binding of SmGrn to any TNR. Interestingly, a weak binding signal with SORT1 (z-score=3) was observed but was below our stringent threshold of z-score >5. Although SORT1 could act as a secondary receptor for SmGrn, its expression in human neutrophils is mostly upregulated in a small subset of less mature proliferative cells: the pre-neutrophils [74]. Our inability to detect binding of the parasite protein on CD177-neutrophils strongly suggests that CD177 is the primary receptor for SmGrn.

In our study, preincubation of neutrophils with SmGrn dampened the inflammatory response to LPS stimulation by reducing CD66b and PRTN3 surface expression as well as reducing PRTN3 proteolytic activity and the secretion of IL-8, and delaying ROS release. In addition, SmGrn-treated neutrophils adopted a more rounded morphology and increased viability, indicative of a more quiescent state. These observations contrast with the activation of human neutrophils observed after crosslinking of CD177 with antibodies [75]. Since CD177 acts as the main anchor for PRTN3 on the neutrophil surface, one possible explanation for the decrease in PRTN3 surface expression and activity is that SmGrn interferes with the formation of the CD177:PRTN3:GPR97 that is required for PAR2 activation and IL-8 release [46]. However, we also observed decreased PRTN3 activity in a CD177^-^ donor, suggesting that SmGrn’s inhibitory effect may also extend beyond the CD177:PRTN3 membrane-anchored complex, possibly by limiting extracellular release of PRTN3 from primary granules. PRTN3 is an important target of anti-neutrophil cytoplasmic antibodies (PRTN3 ANCA), which are responsible for the autoimmune disease granulomatosis with polyangiitis [76]. Interestingly, anti-CD177 Fab fragments that block PRTN3 binding to CD177 have been shown to reduce neutrophil activation in response to PRTN3 ANCA binding [77]. Peptides derived from Granulin proteins have recently been developed for therapeutic purposes. For example, Atsttrin is a peptide derived from subdomains A, C and F of human Progranulin and has been used as an anti-inflammatory drug for its ability to downregulate TNF signalling [78], while Granulin peptides derived from the helminth *Opisthorchis viverrini* have shown wound healing properties through stimulation of fibroblast proliferation [79]. The development of peptides derived from SmGrn that could decrease PRTN3 activity on the neutrophil surface could therefore have beneficial therapeutic effects for the treatment of granulomatosis with polyangiitis.

Finally, neutrophil exposure to SmGrn induced transcriptional changes where almost all differentially expressed genes were upregulated in the presence of the parasite protein, and the functional enrichment was linked to interferon signalling and inflammatory response. Interestingly, amongst the top 20 upregulated transcripts were several genes that have been associated with negative regulation of interferon signalling such as *IDO1* [49], *USP18* [50], *SOCS1* [51], *MT2A* [52] and *NT5C3A* [53]. Transcripts for *IFIT1* and *IL27* were also present in these top 20 transcripts and have been linked to immune suppression by tumour-associated neutrophils [80] and negative regulation of neutrophil function [81], respectively. In accordance with these observations, GO term enrichment analysis of the top 100 transcripts upregulated in SmGrn-treated neutrophils identified IL27-mediated signalling pathway and negative regulation of type II interferon production as the two most enriched biological processes. These observations further support the hypothesis that SmGrn maintains neutrophils in a hyporesponsive state.

The identification of SmGrn as a modulator of human neutrophil activation has uncovered a new mechanism of manipulation of host immunity by *S. mansoni* parasites. By promoting neutrophil quiescence, this may help the parasite establish infection in its human host. Peptides derived from SmGrn may also show therapeutic potential in inflammatory processes where neutrophil activation needs to be dampened.

## Acknowledgements

The study was funded by an MRC fellowship MR/W016397/1 to CC. This UK funded award is carried out in the frame of the Global Health EDCTP3 Joint Undertaking. The authors would like to thank the phlebotomy team and all the blood donors who have contributed to this study; Gavin J Wright for providing access to the collection of plasmids encoding human receptors; Karen Hogg, Andrew Leech and Fabiano Pais from the Bioscience Technology Facility at the University of York for technical assistance; Drs Jarrod Shilts, James Hewitson, Dave Boucher and Borko Amulic for helpful comments and advice. This study is dedicated to the memory of our friend and colleague, Dr Kelly Lee.

## Authors’ contributions

MM performed the flow cytometry and functional analysis of human neutrophils, KL did the biochemical validation and domain-mapping of the interaction, NMS performed the large-scale protein:protein interaction assay, CC conceived the study, performed the large-scale protein:protein interaction assay and identified the interaction. MM and CC wrote the manuscript with contributions from all the authors. The authors declare no competing interests.

---

**Supplementary figure 1.**
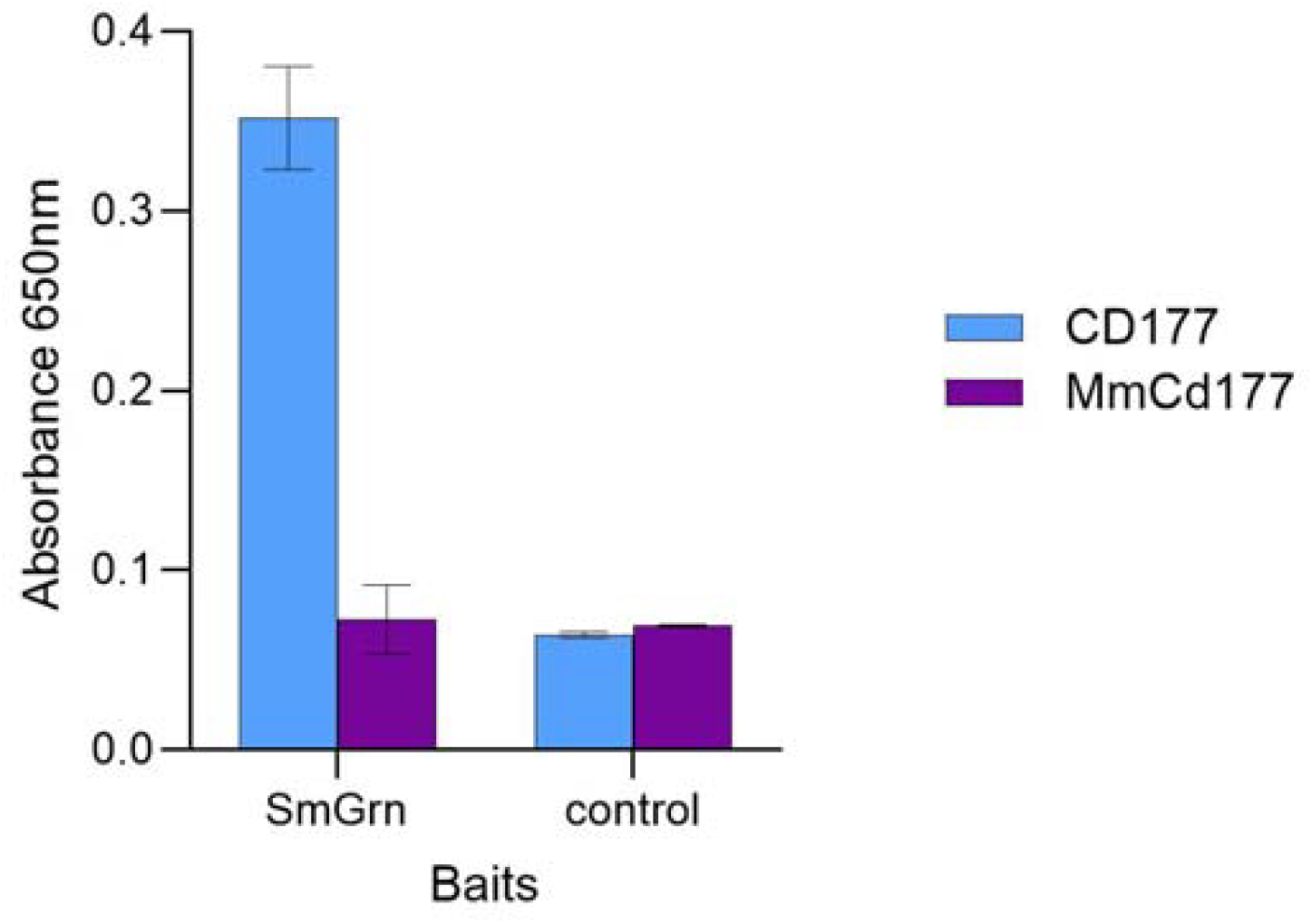
Murine Cd177 does not interact with SmGrn. SmGrn or the tag only control were immobilised as baits on a streptavidin-coated plate and incubated in the presence of either human CD177 (CD177, blue) or mouse Cd177 (MmCd177, purple). Interaction was detected only between SmGrn and the human receptor. Bars represent mean +/-SD, n=3 technical replicates. One representative of two biological replicates.

**Supplementary figure 2.**
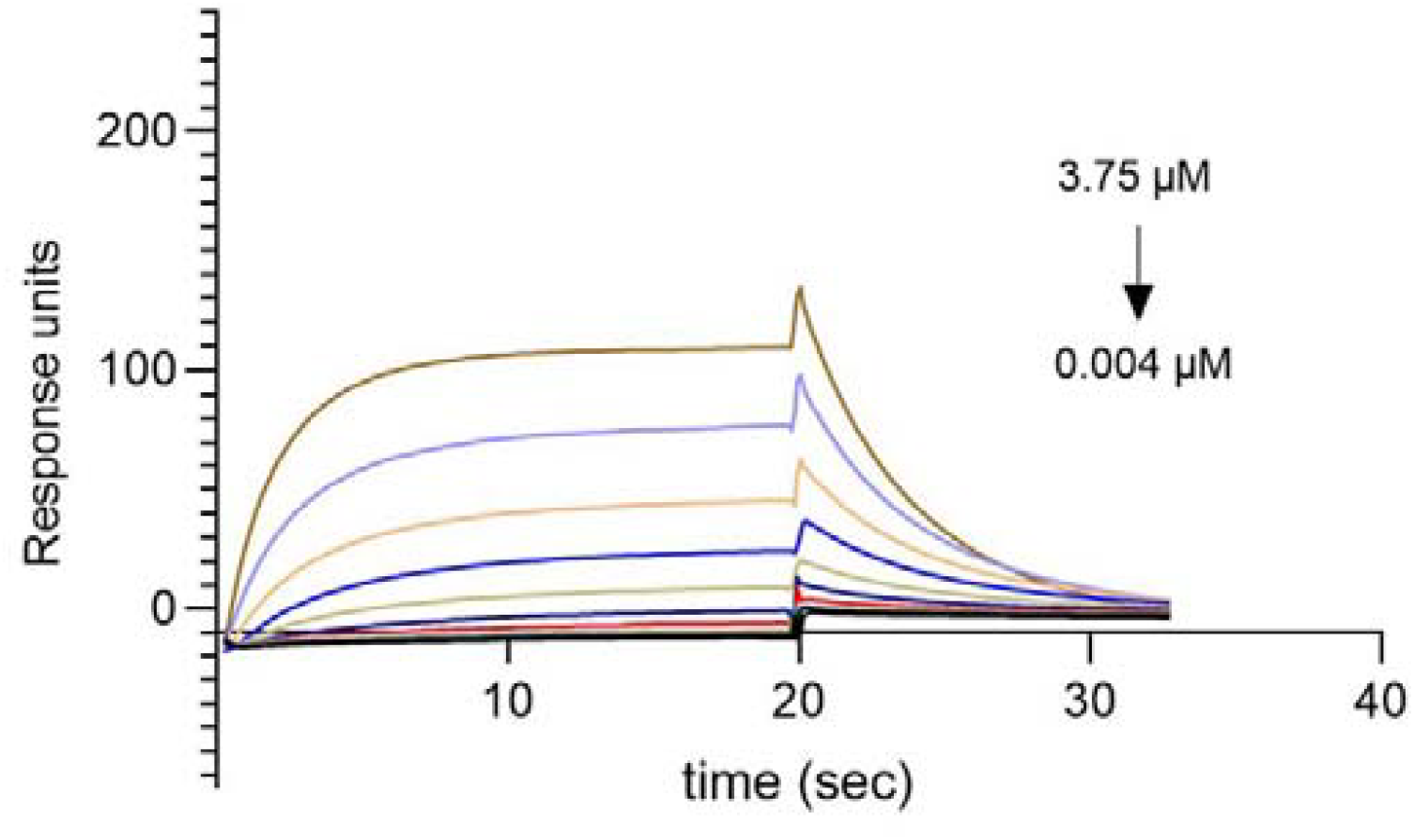
Kinetics analysis of SmGrn binding to CD177. Increasing concentrations of SmGrn were injected at high flow rate over a chip containing the CD177 ligand. Kinetics analysis assuming a 1:1 binding model produced an association rate constant *k*_*a*_ = (1.76 ± 0.08) x 10^5^ M^-1^ s^-1^ and a dissociation rate constant *k*_*d*_ = 0.22 ± 0.01 s^-1^ resulting in a *K*_*D calc*_ = 1.2 μM.

**Supplementary figure 3.**
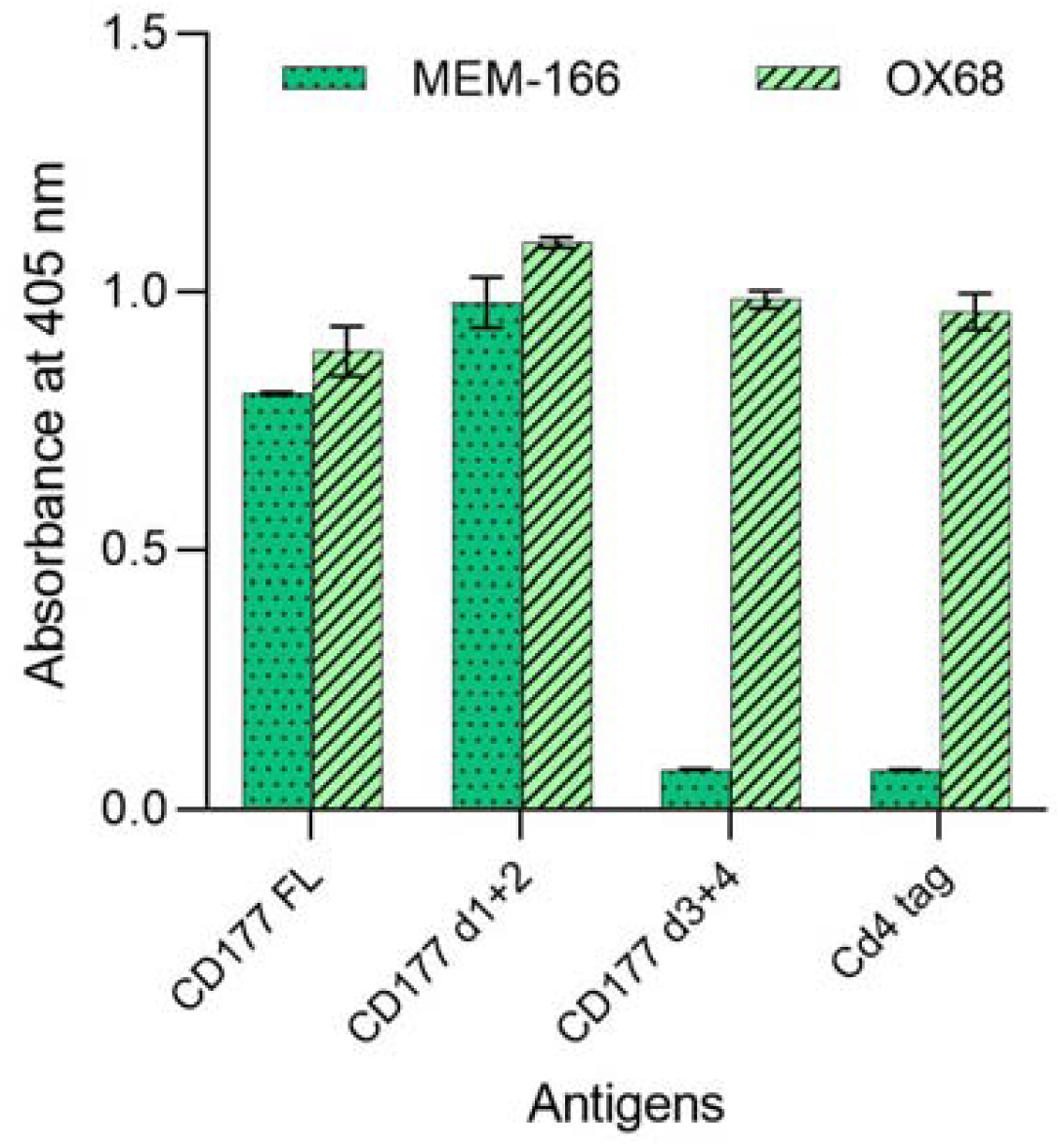
The MEM-166 antibody binds the N-terminal subdomain of CD177. Biotinylated proteins corresponding to the Cd4-tagged full-length ectodomain of CD177 (CD177 FL), its N-terminal (CD177 d1+2) or C-terminal (CD177 d3+4) subdomains or the Cd4 tag alone were immobilised of a streptavidin-coated plate and incubated with either the anti-CD177 antibody MEM-166 or the anti-Cd4 tag antibody OX68. Binding of the primary antibodies was revealed by an alkaline-phosphatase-conjugated secondary antibody and a phosphatase substrate, demonstrating binding of MEM-166 to the amino-terminal domain of CD177.

**Supplementary figure 4.**
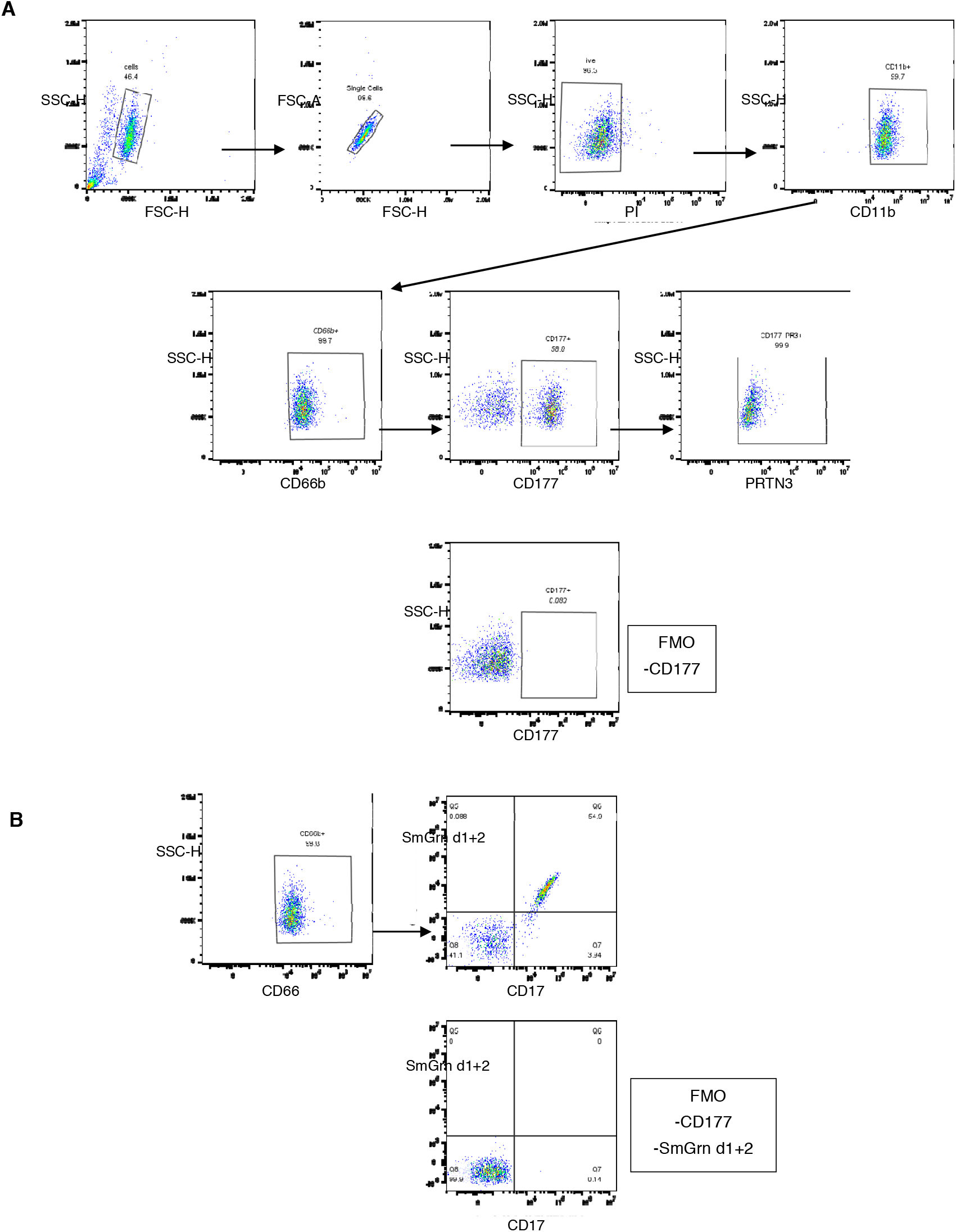
Gating strategies used for the staining of human neutrophils. **A**. Forward scatter (FSC) and side scatter (SSC) were used to gate on cells and exclude cellular debris, single cells were then separated from doublet by FSC analysis and PI-positive apoptotic cells excluded from the analysis. CD11b and CD66b were used to identify mature neutrophils, which could be separated into CD177^+^ and CD177^-^ subpopulations. Only the CD177+ population showed surface staining for PRTN3. **B**. The same gating strategy was used to isolate CD66b^+^ neutrophils and the level of CD177 staining compared to the level of SmGrn d1+2 binding to the neutrophil surface. Fluorescence minus one (FMO) controls are shown on the bottom row for each gating procedure.

**Supplementary figure 5.**
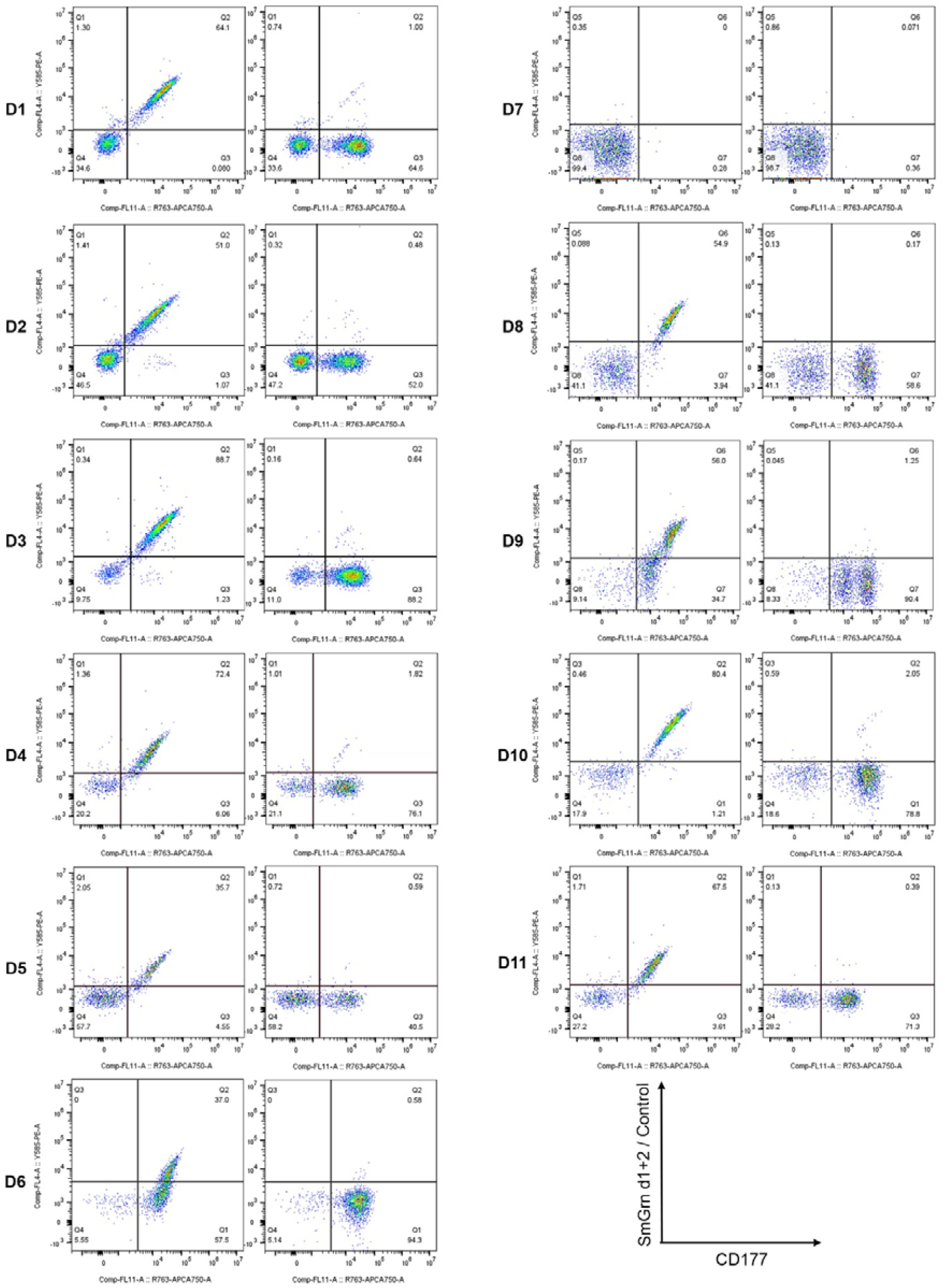
SmGrn binds the surface of CD177^+^ human neutrophils. Flow cytometry dot plots of blood-isolated neutrophils from 11 donors with different proportions of CD177^+^ neutrophils. Cells were stained with an anti-CD177 antibody and either biotinylated SmGrn d1+2 (left panel) or a Cd4 d3+4 tag only as a control (right panel) and detected with streptavidin-PE.

**Supplementary figure 6.**
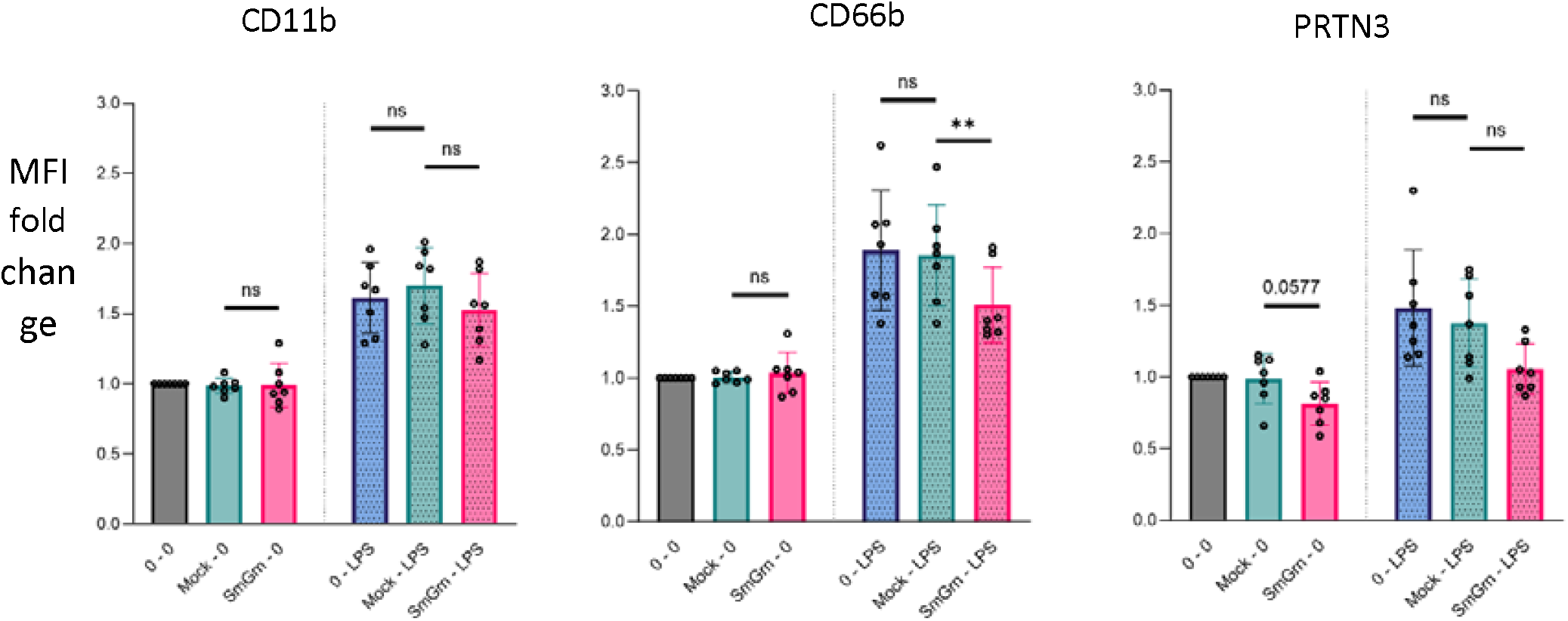
MFI fold changes of CD11b, CD66b and PRTN3 for all experimental conditions. Each donor is represented by a dot and the MFI corresponding to untreated-unstimulated (0-0) conditions was normalised to 1. T-test for unstimulated groups, and one-way ANOVA (followed by Dunnett’s multiple comparisons test) for LPS-stimulated groups. Statistical significance is indicated by ** P<0.01.

**Supplementary figure 7.**
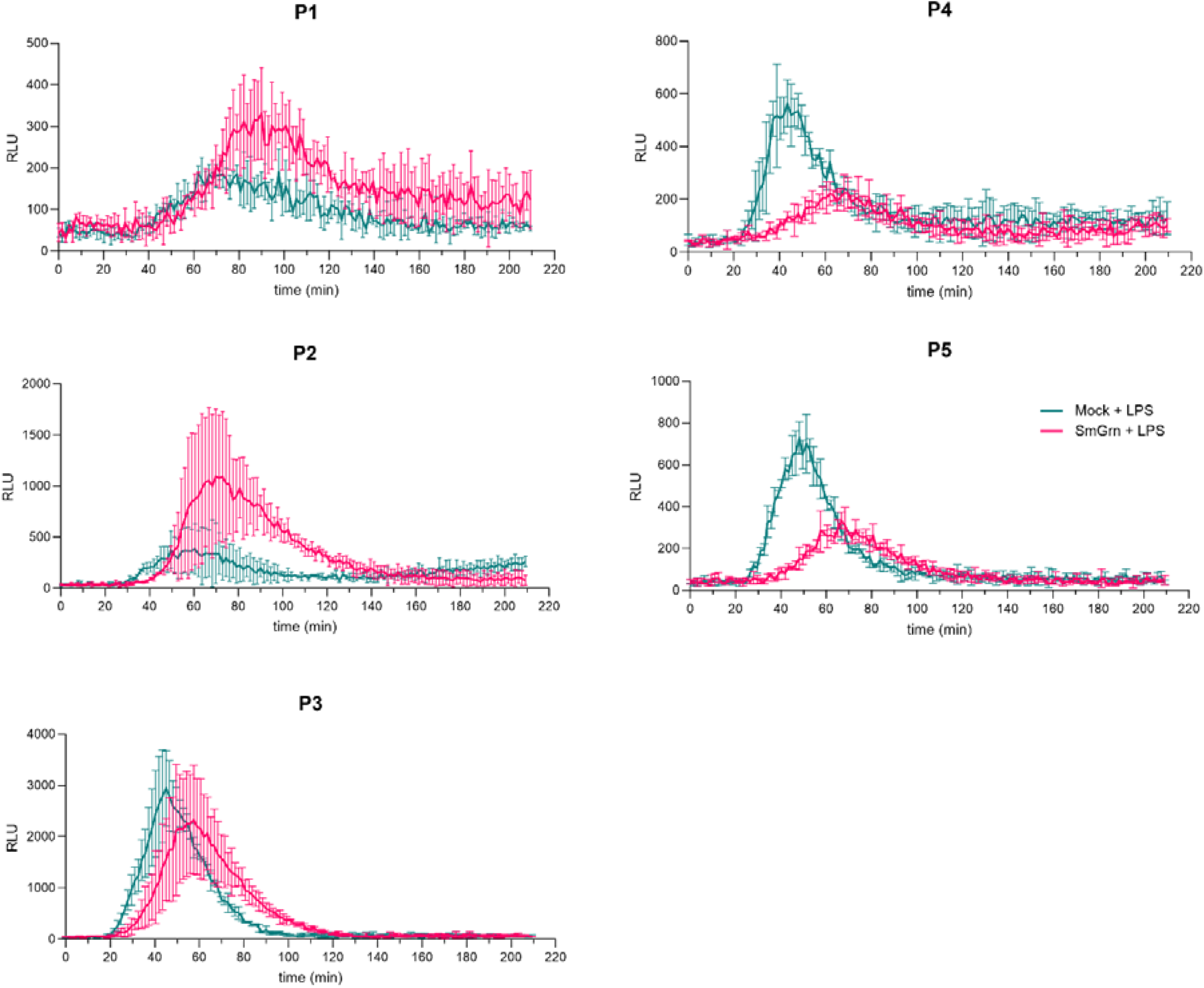
Reactive oxygen species release is delayed in neutrophils pre-incubated with SmGrn. ROS release was assessed by median luminescence over a period of three hours from the neutrophils of five different participants (P1 to P5) after pre-incubation with either mock-transfectant (green) or SmGrn (pink). Delay in ROS release from neutrophils pre-incubated with SmGrn was observed in all donors. n = 5 different donors from 2 independent experiments with fractions of CD177^+^ neutrophils ranging from 45% to 93%. Error bar = SD from 3 technical replicates.

**Supplementary figure 8.**
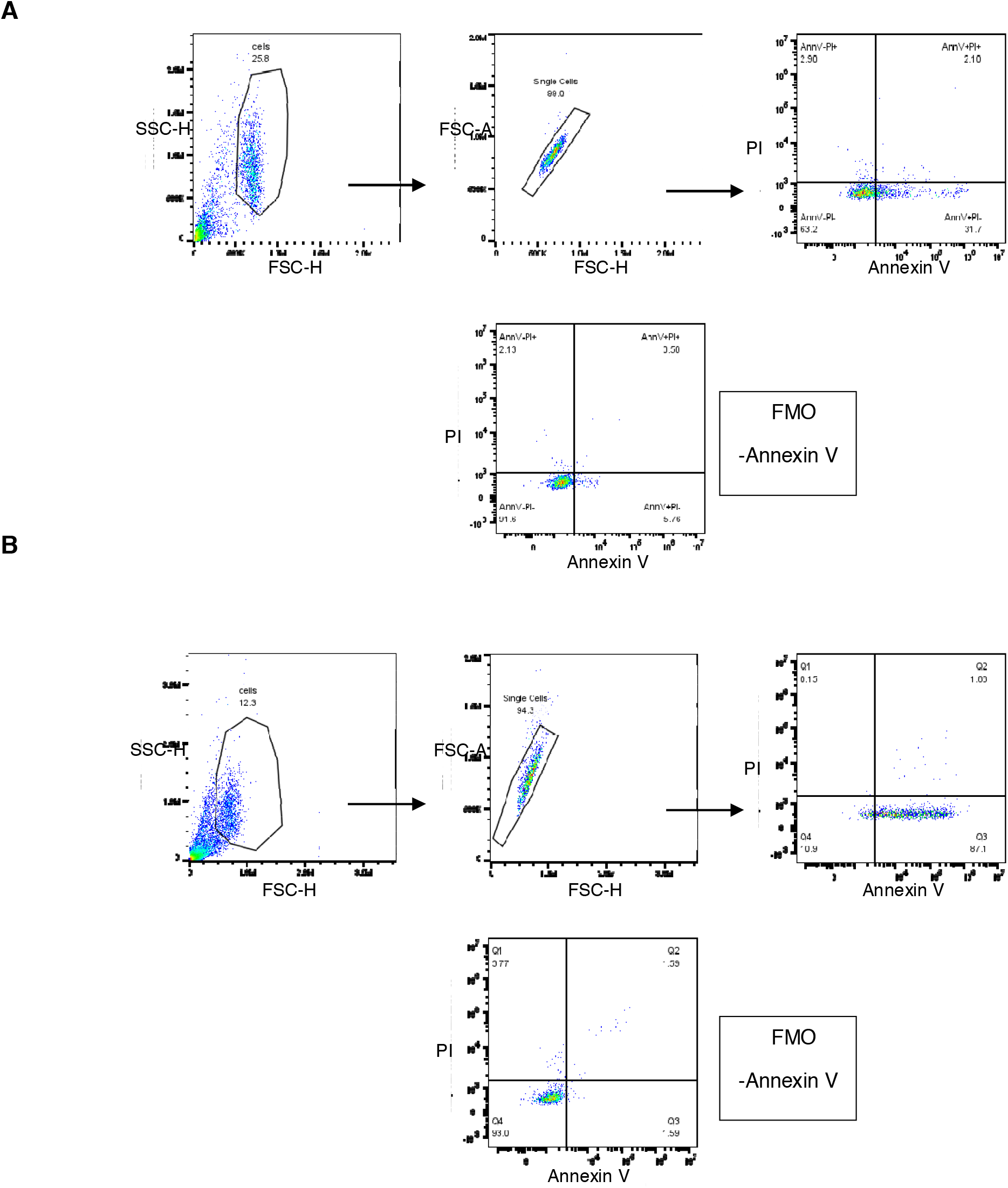
Gating strategy used for the staining of apoptotic neutrophils. **A**. Forward scatter (FSC) and side scatter (SSC) were used to gate on cells and exclude cellular debris; single cells were then separated from doublets by FSC analysis. Early and late apoptotic cells were identified by Annexin V and PI stainings, respectively. **B**. The same gating strategy was used to evaluate the effect of staurosprorine treatment on isolated neutrophils. Fluorescence minus one (FMO) controls are shown on the bottom row for each gating procedure.

**Supplementary figure 9.**
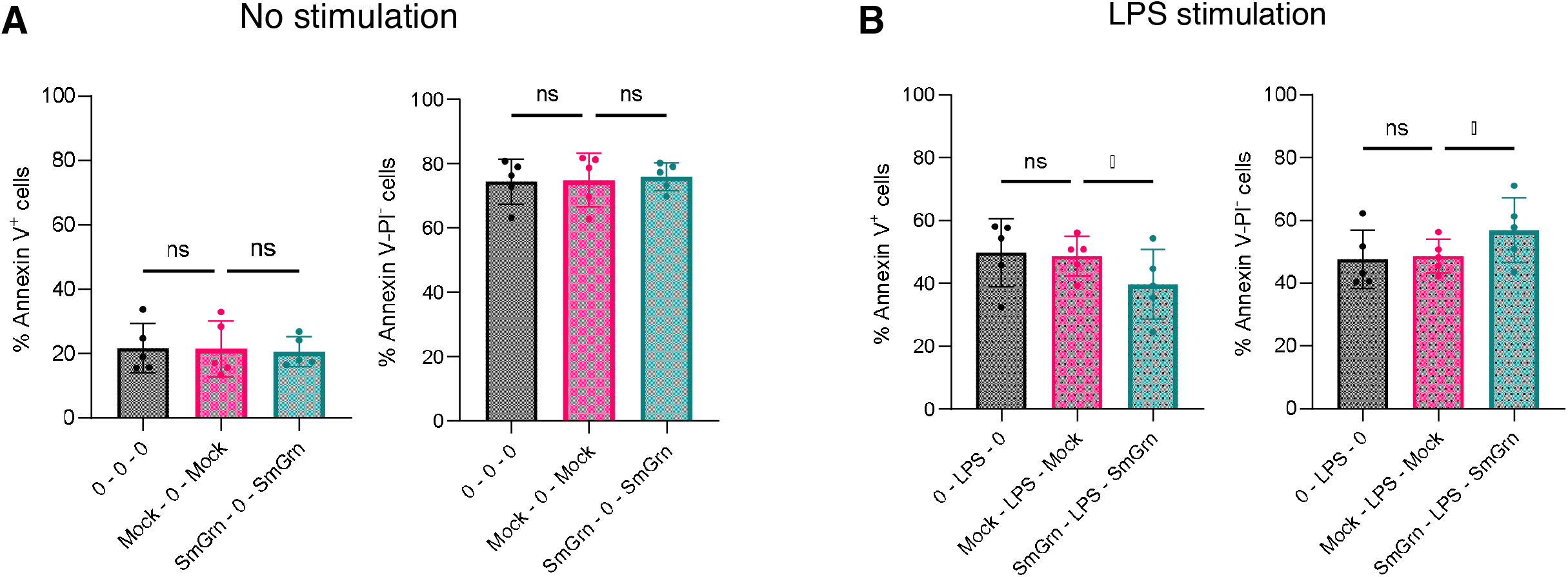
Preincubation of human neutrophils with SmGrn increases their viability. **A**. Percentages of early apoptotic (Annexin V^+^) neutrophils (left) and viable (Annexin V-PI^-^) neutrophils (right) in the absence of stimulation. **B**. Percentages of early apoptotic (Annexin V^+^) neutrophils (left) and viable (Annexin V-PI^-^) neutrophils (right) following stimulation with LPS. One-way ANOVA comparison (followed by Dunnett’s multiple comparisons test). Statistical significance is indicated by * P<0.05.

